# Post-Synaptic Density Proteins in Oligodendrocytes are Required for Activity-Dependent Myelin Sheath Growth

**DOI:** 10.1101/2025.02.10.637467

**Authors:** MA Masson, M Graciarena, M Porte, B Nait Oumesmar

## Abstract

Compelling evidence demonstrates a functional link between neuronal activity and myelination, highlighting the vital importance of axon-oligodendrocyte crosstalk in myelin physiology and function. However, how neuronal activity is relayed to oligodendroglia to regulate myelin formation remains not fully understood. Here, we aimed to characterize how that myelination is regulated by glutamate vesicular release in zebrafish spinal cord. We compared oligodendrocyte precursor cells (OPCs) and myelinating oligodendrocytes (mOLs) for their close apposition with pre-synaptic boutons and found that these are increased in number on mOLs during myelin internode elongation. Consistently, mOLs show more pre-synaptic boutons during myelin internode elongation compared to OPCs. In addition, we also found that oligodendroglial cells express the post-synaptic density protein 95 (PSD-95) along punctated domains, regardless of their differentiation stage. Genetically targeted PSD-95-GFP expression in oligodendroglia revealed post-synaptic-like domains along their processes and sheaths, which are contacted by axonal pre-synaptic varicosities. These contacts are increased in mOLs. Importantly, CRISPR-Cas9 mediated deletion of *dlg4* in oligodendroglia impairs myelin sheath growth*, in vivo*. Overall, our data indicate that PSD-95 is a key component of axons to oligodendrocytes neurotransmission that regulates myelin sheath growth.

**HIGHLIGHTS:**

- Glutamate vesicular release is required for myelination
- Axon-oligodendroglia connectivity increases with oligodendrocyte maturation
- Oligodendrocytes express the post-synaptic density protein 95
- *Dlg4* loss-of-function in oligodendroglia impedes myelin sheath growth

**GRAPHICAL ABSTRACT:** 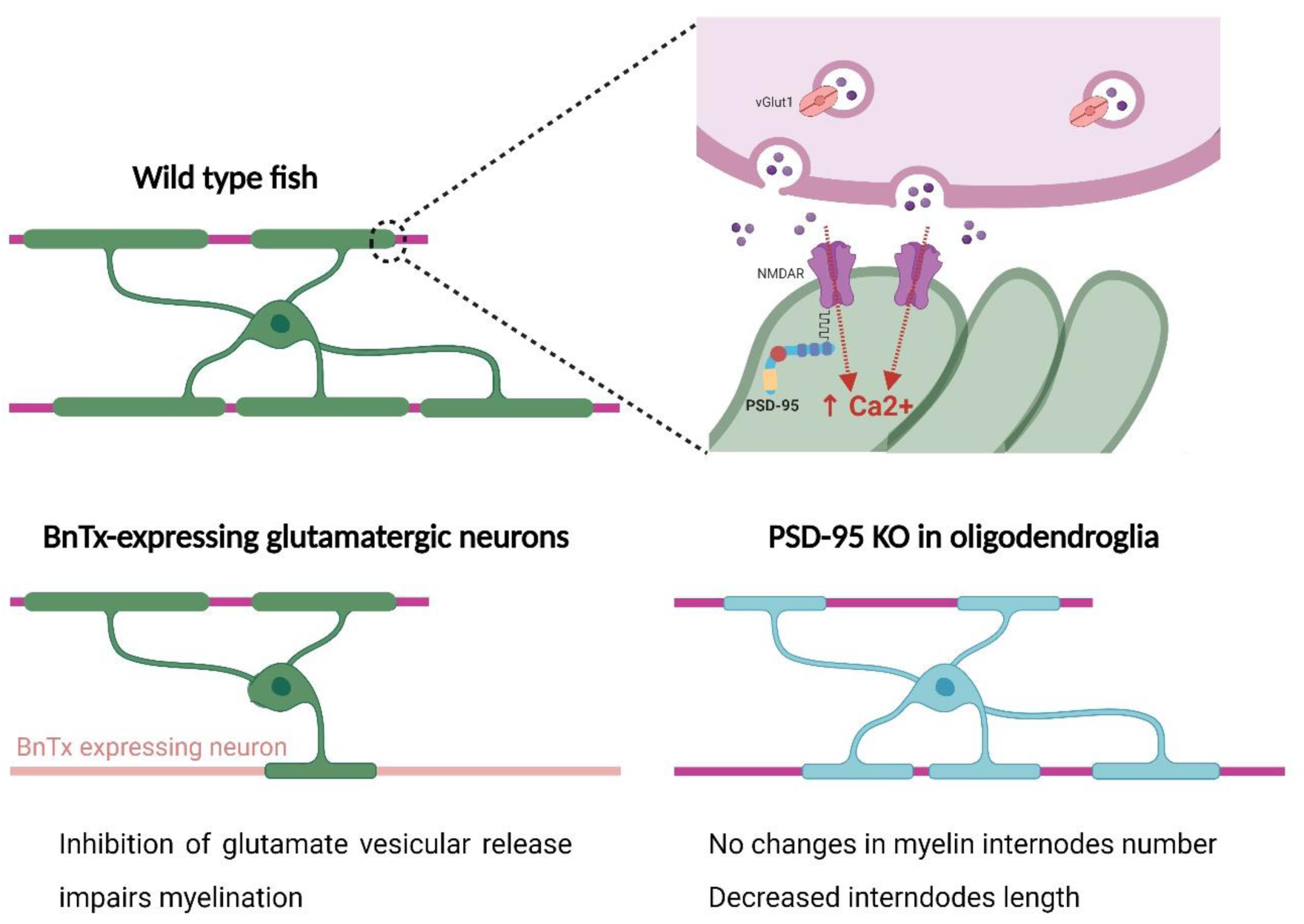

## Introduction

Myelin is a multilamellar lipid rich structure insulating axons and allowing the saltatory conduction of action potentials. In the central nervous system (CNS), oligodendrocytes (OLs) preferentially myelinate electrically active axons ^1–5^, supporting the notion that neuronal activity regulates myelination. In turn, adult myelination regulates complex behaviors ^6–8^. Therefore, changes in myelin microstructure sense and affect the functional connectivity of neural circuits, underlying vital neurological functions. OLs derive from oligodendrocyte precursor cells (OPCs), which constitute the major cellular source for myelin regeneration ^9^. OPCs receive AMPA- and NMDA-mediated synaptic inputs from neuronal fibers throughout the CNS (^10–12^ and others), with AMPA receptors exerting a positive role on myelination ^13^. NMDA receptors are also expressed in oligodendroglial cells including myelinating oligodendrocytes (mOLs) ^14,15^ but their role in myelination is still controversial ^16–19^. Importantly, these synapses are generally thought to be restricted to the OPC stage and rapidly lost upon differentiation into mOLs ^20,21^. However, recent evidence indicates that glutamate induces calcium transients in myelin sheaths ^22–24^. In addition, electrical activity affects myelin retraction and stabilization in nascent sheaths rather than myelination onset ^4,5,24^.

The nature and extent of connectivity between axons and oligodendroglial cells during lineage progression and myelination remain poorly characterized. Here, we show that vesicular release of glutamate plays a key role in myelin sheath growth in zebrafish larvae. Among oligodendroglial cells, mOLs undergoing sheath extension bear the highest extent of pre-synaptic vesicle appositions. Interestingly, we showed that PSD-95, a major scaffolding PDZ protein of glutamatergic post-synaptic densities in neurons, is expressed by oligodendroglia at all differentiation stages. However, PSD-95 clusters, as a counterpart of pre-synaptic boutons, are highest in mOLs undergoing sheath extension. Remarkably, oligodendroglia cell-specific knock-out (KO) of *dlg4* showed impaired myelin sheath elongation. Taken together, our data reveal an increasing connectivity throughout oligodendroglial lineage progression, highlighting mOLs as a cellular target for neuronal activity.

## Results

### Myelinating oligodendrocytes establish synaptic-like connectivity with axons

Broad evidence showed that electrical activity regulates myelination, though synaptic vesicular release of neurotransmitters along axons ^2,25^. However, the mode of communication and the extent of connectivity of mOLs with axons remain still poorly understood. To decipher the modes of interaction between OLs and axons, we assessed the impact of glutamate vesicular release on myelination *in vivo*. To do so, we generated the *Tg(vGlut2a:Gal4;UAS:BnTx/LC:GFP::sox10:mRFP)* ^26,27^ zebrafish line, where glutamatergic neurons visualized with GFP have their vesicular release impaired by botulinum toxin (BnTx), while *sox10-mRFP* labels the oligodendroglial population (Figure 1A). At 5 days post-fertilization (dpf), we observed a marked decrease of mRFP-positive myelin area in BnTx-expressing larvae (Figure 1C, D) with respect to controls (Figure 1B). Myelination impairment in BnTx-expressing larvae was further confirmed by electron microscopy (EM) (Figure 1E, F) and quantification of myelinated axons in the ventral spinal cord, which was significantly reduced in BnTx-expressing zebrafish larvae (Figure 1G). To exclude any potential effects of BnTx on neuronal and oligodendroglial cell proliferation or death, we also quantified these cell types in control and BnTx-expressing larvae, at 5 dpf. The number of HuC-positive neurons and Olig2-DsRed oligodendroglial cells did not reveal any significant changes in BnTx-expressing zebrafish, indicating that myelination impairment was not accounted to axonal loss nor to oligodendroglial cell death (data not shown).

**Figure 1:**
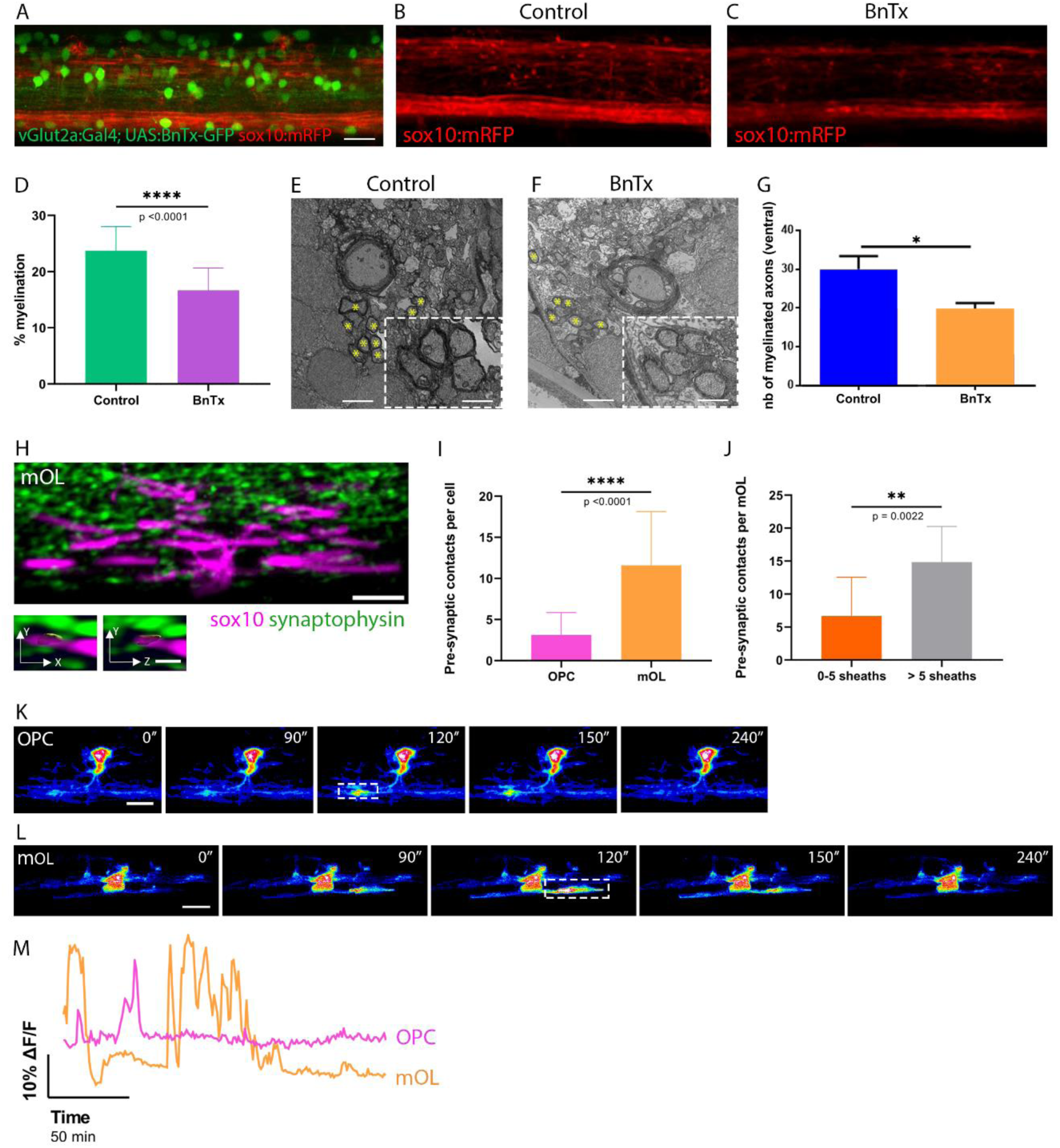
Myelinating oligodendrocytes establish synaptic-like connectivity with axons. A. Labelling pattern of zebrafish *vGlut:Gal4; UAS:BnTx-GFP::sox10:mRFP* reporter line. Scale bar: 20 µm. B-C. Comparative view of sox10 labeling in control (B) and BnTx-positive (C) zebrafish in the larval spinal cord at 5 dpf. D. Percentage of myelinated axons in the ventral region of hemispinal cords. Student’s t test; N= 14 controls vs 15 BnTx larvae. E-F. Electron microscopy analysis of myelination in control (E) and BnTx-positive (F) zebrafish. Myelinated axons are highlighted with yellow stars. Scale bar: 2 µm. Inset: 1 µm. G. Myelinated axon number in the ventral region of hemispinal cords. Student’s t test. H. Analysis of synaptic vesicle juxtaposition to mOL. Insets show axon-OL contacts with pre-synaptic vesicles. Scale bars: 10 µm. Insets: 2 µm. I. Number of synaptic-like boutons on OPCs and mOLs. Student’s t test; N= 16 vs N=24 cells respectively. J. Number of contacts on mOL that bear 0-5 sheaths and >5 sheaths. Mann-Whitney test; N= 8 versus 12 cells. K-L. Still pictures of live imaging in (K) an OPC and (L) a mOL showing calcium transients in processes and myelin. M. Pattern of spontaneous calcium transients in OPCs and mOLs. Analyzed regions are shown in white in (K) and (L). Scale bars: 10 µm.

Next, we investigated at which stage of differentiation are oligodendroglial cells most receptive to neuronal activity. We analyzed synaptic vesicle-mediated apposition between axons and oligodendroglial cells by generating *Tg(Huc:Gal4; UAS:synaptophysin:GFP::sox10:mRFP)* ^27,28^ zebrafish, where pre-synaptic vesicles were visualized by GFP, and subsequently chimera fish generated by cell transplantation from the *sox10:mRFP* line into the *Huc:Gal4; UAS:synaptophysin:GFP* line. We observed synaptic vesicles closely apposed to oligodendroglial cells based on the partial overlap between synaptophysin:GFP and sox10:mRFP fluorescence (Figure 1H). Quantification of pre-synaptic inputs onto OPCs and mOLs revealed a higher number of pre-synaptic boutons on mOLs (Figure 1I). Among mOLs, it is noteworthy that the number of pre-synaptic boutons has a wider range of distribution, suggesting that this might be correlated to different stages of the myelination process. In the zebrafish, OLs take up to 5 hours to establish new myelin sheaths and mostly sheath elongation occurs after this time window ^29^. The final number of sheaths per OL is typically between 5 and 15 ^30^. Based on these data, we classified mOLs according to their sheath number: mOLs with less than 5 sheaths that should be undergoing new sheaths generation and mOLs with more than 5 sheaths, that are mostly undergoing myelin sheath elongation. When comparing the number of pre-synaptic boutons, mOLs with more 5 sheaths had significantly more pre-synaptic inputs (Figure 1J). Overall, our data indicate mOLs are contacted by axons through synaptic-like communications, which increase in number during myelin sheath elongation.

For a functional correlate of pre-synaptic inputs on oligodendroglial cells, we tested OPCs and mOLs for spontaneous calcium transients by injecting a *sox10:GCaMP6* construct into wild type one-cell stage embryos and recorded calcium transients in 3 dpf larva spinal cords. As previously reported, both OPCs and mOLs display calcium transients (Figure 1K, L; Videos S1 and S2) in their processes and sheaths ^23,24^. Consistent with their higher number of pre-synaptic inputs, mOLs display calcium transients with a higher frequency and amplitude with respect to those of OPCs (Figure 1M). Thus, the extent of axonal communications with oligodendroglial cells is highest during the active period of myelination.

### PSD-95 post-synaptic densities increase in myelinating oligodendrocytes

One of main structural feature of chemical synapses is the presence of a post-synaptic density (PSD). Hence, we searched for the expression of the post-synaptic density-95 (PSD-95) in oligodendroglial cells, PSD-95 is a major scaffolding protein that has been mainly described at the post-synaptic density of glutamatergic synapses in neurons ^31^. However, PSD-95 expression in glia has been poorly studied. Interestingly, we found punctuated PSD-95 expression along *Tg(Olig1:KalTA4, UAS:mCherry)* zebrafish spinal cord, at 5 dpf. Olig1-expressing OPCs and OLs expressed PSD-95 on their cell soma and processes as shown by immunohistochemistry (data not shown). As expected, it is worth noting that PSD-95 was also broadly expressed in neurons. Using IMARIS 3D reconstruction of Olig1-expressing cells and PSD-95 puncta, we selected PSD-95 puncta located on oligodendroglia soma and processes (Figure 2A, B). To determine whether oligodendroglial PSD-95 is targeted to specific contact sites with axons, we generated a tol2-flanked *sox10:PSD-95-GFP* expression vector and injected it into one-cell stage eggs of the *Tg(vGlut:Gal4;UAS:mRFP)* ^26^ line. This resulted in mosaic oligodendroglial cells labeled with PSD-95-GFP fusion protein at their target sites on glutamatergic axons stained with mRFP (Figure 2C). Using *in vivo* live imaging in the larval zebrafish spinal cord, we quantified the number of juxtapositions between glutamatergic axonal varicosities and PSD-95-GFP domains on oligodendroglial cells, at 3dpf (Figure 2C, insets). Additionally, we corroborated that axonal varicosities in *vGlut:Gal4;UAS:mRFP* line reliably represent sites of pre-synaptic vesicle accumulation, as they co-localize with synaptophysin-GFP puncta (Figure 2D). Importantly, we found that the number of glutamatergic communications increased on mOLs (Figure 2E). We next categorized mOLs according to their sheath number, as described previously ^29^, and compared the number of glutamatergic contacts onto mOLs that had less or more than 5 sheaths. Consistent with the pre-synaptic bouton analysis, we found that post-synaptic densities increased significantly on mOLs with more than 5 sheaths, mostly undergoing myelin internode elongation (Figure 2F). Altogether, our findings clearly demonstrate that glutamatergic synaptic-like communications are maintained throughout the oligodendroglial lineage and increase in number at the myelinating stage. As myelination is actively shaped by glutamate vesicular release, our results point to mOLs as the main cell target of neuronal activity.

**Figure 2:**
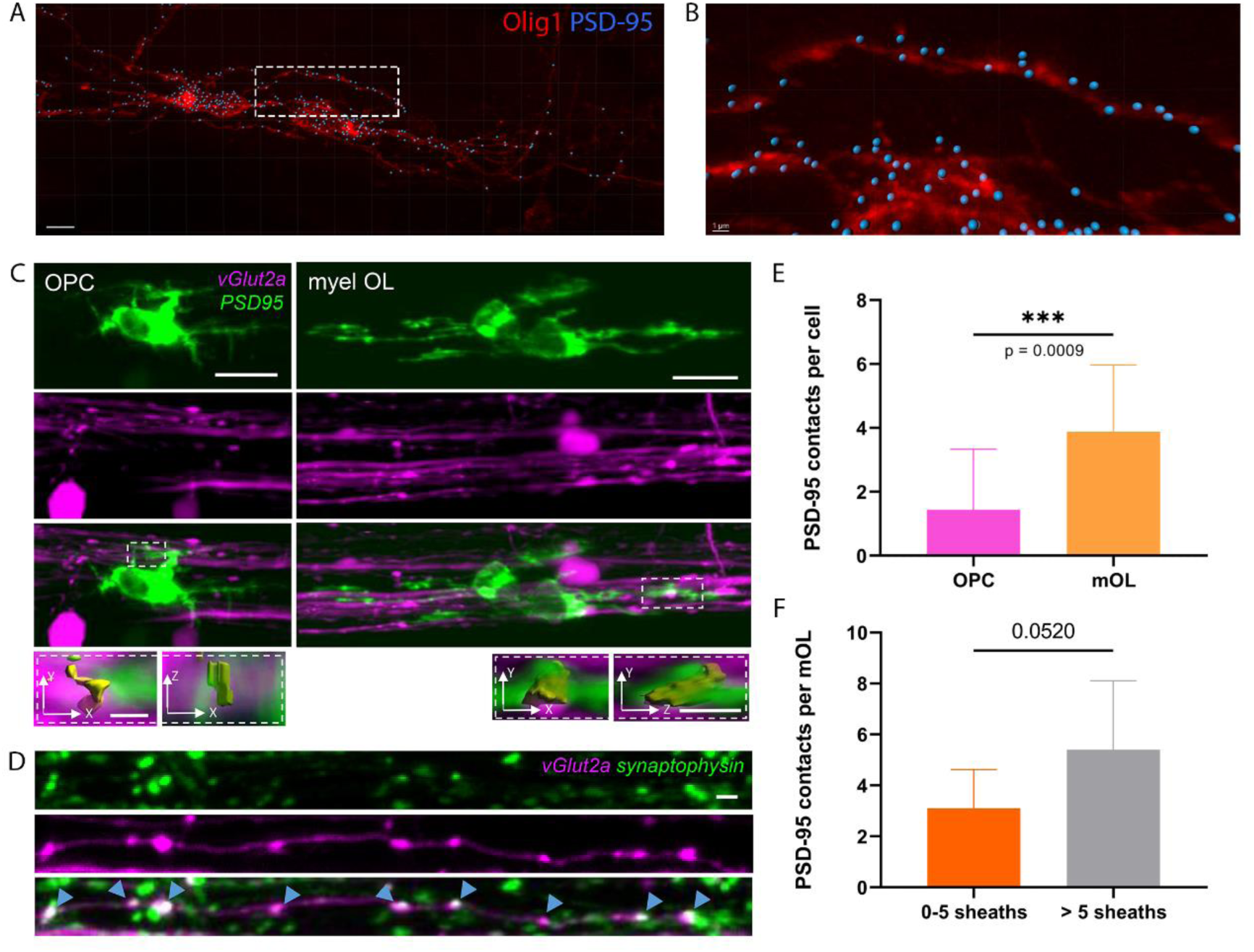
PSD-95 post-synaptic densities in oligodendroglia increase during active myelination. A. PSD-95+ puncta (blue) located on Olig1+ oligodendroglia soma and processes (red) in the zebrafish spinal cord at 5 dpf. Cell surfaces and puncta were reconstructed with IMARIS. Scale bar: 7 µm. B. Inset of the oligodendroglial process in (A). Scale bar: 1 µm. C. Targeted expression of PSD-95-GFP in OPCs (left panel) and mOLs (right panel) of the zebrafish spinal cord. Insets show the contacts based on partial overlapping fluorescence. Scale bars: 10 µm. Insets: 2 µm. D. Co-localization between varicosities of glutamatergic axons *(vGlut:Gal4;UAS:mRFP)* and pre-synaptic vesicles (*HuC:Gal4;UAS:synaptophysin-GFP*, arrowheads). Scale bar: 2 µm. E. Quantification of synaptic-like contacts in OPCs and mOLs. Mann-Whitney test; N=16 cells for each group. F. Number of synaptic contacts in mOLs that bear 0 to 5 sheaths and >5 sheaths. Student’s t test; N=10 versus 5 cells, respectively.

### PSD-95 loss-of-function in oligodendroglia impairs myelin sheath elongation

In zebrafish, the *dlg4* gene, encoding for PSD-95, is duplicated in two isoforms *dlg4a* and *dlg4b*, both expressed in oligodendroglial cells ^32^. To determine the functional role of PSD-95 in oligodendroglia, we used a cell-specific CRISPR-Cas9 mediated loss-of-function approach, based on the Gal4-UAS system and generated a Tol2 expression vector encoding a specific *dlg4a* single guide RNA (sgRNA) under the control of the U6 promotor, the Cas9 endonuclease and the mCherry fluorescent reporter under the control of UAS cassette (Figure 3A). As the *dlg4a* gene possesses four variants, we targeted the first common exon – the exon 5 – of these variants. Using the CRISPR tool, we based our sgRNA selection choice on two algorithms: the MIT specificity score (>90) ^33^ and Cas9 predicted efficiency (>50) ^34^. We also considered the number of off-targets that must be null or minimal. We selected sgRNA that have no potential off-targets for sequences with less than three base pairs mismatches.

**Figure 3:**
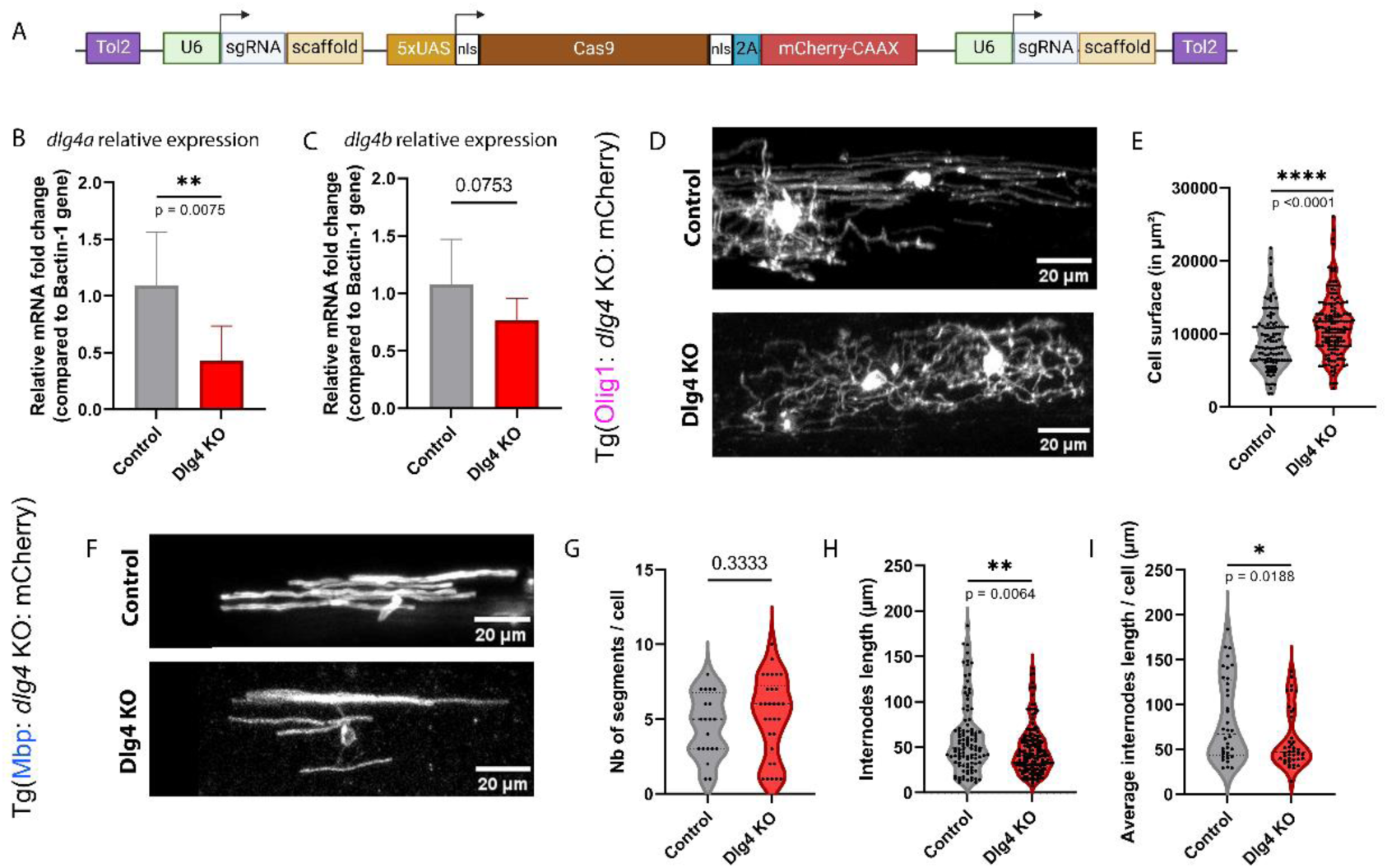
CRISPR-mediated PSD-95 loss-of-function in oligodendroglia impairs myelin sheath elongation. A. Representation of the plasmid micro-injected into one-cell stage of *Tg(s1020t:Gal4)* (B, C), *Tg(Olig1:KalTA4)* (D, E) or *Tg(Mbp:Gal4)* (F-I) to induce cell-specific dlg4a loss-of-function with the Cas9 endonuclease. B-C. Quantitative RT-PCR, on 1020+ mCherry+ whole larvae, of (B) *dlg4a* and (C) *dlg4b* gene expression at 5dpf. Student’s t test; N=4 versus 7 groups of animals, each group containing RNA from 20 dissociated fish. Data are represented as mean ± SD. D. Olig1+ cells in control and *dlg4* KO spinal cords at 5 dpf exhibit distinct morphologies. In *dlg4a* KO, Olig1+ cells are more ramified than in the control. Mature myelinating OLs were mainly observed in control animals. E. Quantification of the oligodendroglial cell surface in control and *dlg4a* KO animals with IMARIS. Mann-Whitney test; N=117 vs 162 cells, respectively in control and KO fish. F. MBP+ cells with myelin segments in control and KO spinal cords at 5 dpf. G. Quantification of the number of myelin segments / cell. Mann-Whitney test; N=20 vs 30 cells, respectively. H. Quantification the internode length. Mann-Whitney test; N=107 vs 165 internodes. I. Quantification of the internode length per cell. Mann-Whitney test; N=36 vs 41 cells.

We next validated our sgRNA and construct under the *s1020t:Gal4* promoter ^35^, as it is strongly expressed as soon as 1 dpf. In mCherry+ whole larvae, we observed a 57% decrease in the *dlg4a* relative expression by RT-qPCR (Figure 3B) and no change in *dlg4b* relative expression (Figure 3C). The construct was then microinjected into *Tg(Olig1:KalTA4)* ^32^ zebrafish eggs and recombined fish were selected based on the expression of the mCherry fluorescence. At 5 dpf, we observed that *dlg4a* knock-out (KO) cells exhibit more ramifications than control cells (Figure 3D, Videos S3 and S4) and display the morphology of immature OLs. Using 3D reconstruction of mCherry-expressing oligodendroglial cells, we demonstrated that *dlg4a* KO cells had a larger cell surface than controls (Figure 3E), indicating that PSD-95 loss-of-function in oligodendroglia impedes myelination. We next assessed the myelination capacity of *dlg4a* KO cells using the *Tg(Mbp:Gal4)* ^30^ line for a specific deletion of PSD-95 in mature OLs and quantified the number of internodes per cells and the internode length. A 5 dpf, during the active period of myelination, Mbp+ mOLs in both control and *dlg4a* KO animals had the same number of myelin segments per cell (Figure 3F, G). However, in *dlg4a* KO, myelin internodes were significantly shorter (Figure 3H, I), demonstrating that oligodendroglial PSD-95 plays a critical role in myelin internode growth.

## Discussion

In the present study, we showed that inhibition of glutamate vesicular release hampers CNS myelination. In line with this, electrical activity is known to bias OL preference for actively electrically axons ^4^ and affects myelin sheath number of single OLs ^5^. Recently, it has been shown that glutamate vesicular release occurs at heminodes of myelin segments, where vGlut1 and neurofascin proteins also co-localized, and reinforces nascent sheaths elongation ^25^. Moreover, translation of mRNA encoding myelin proteins is regulated by NMDA receptors on oligodendroglial cell processes ^2,36^. Thus, silent glutamatergic axons might be selectively unmyelinated in the BnTx-expressing fish due to a failure in myelin mRNA translation at specific ensheathment sites. Interestingly, specific subpopulations among glutamatergic neurons exhibit diversity in their dependence on synaptic vesicle release for myelination ^25,37^. As the overall reduction of myelination upon inhibition of synaptic glutamate vesicular release is broad enough, one could speculate that neuronal activity shapes myelin formation preferentially on glutamatergic axons; this hypothesis awaits further exploration.

We also found that the number of pre-synaptic boutons is higher once myelination has started. Consistently, both OPCs and mOLs display spontaneous calcium currents ^15,23,24,38^, with increased frequency and amplitude in mOLs. Thus, axonal signaling to oligodendroglial cells may be more prominent during active myelination. In agreement with this hypothesis, synaptic vesicles accumulate at myelin ensheathment sites ^4,39^. However, in our study, we also observed axonal connectivity on OPCs that is supported by broad evidence on axon-OPC synapses ^10,11,40^. mOLs composed of more than 5 myelin sheaths, which are mainly undoing myelin sheath elongation, bared a higher number of contacts with axons at specific sites of synaptic vesicle accumulation. Hence, the highest extent of synaptic vesicle mediated communications between axons and mOLs may occur preferentially during myelin sheath growth. A comprehensive analysis of how pre-synaptic vesicles are transported to and accumulate at restricted axoplasm domains during myelin sheath formation could greatly improve our understanding of activity-dependent myelin formation. In line with this, by using the genetically encoded sensor of vesicle exocytosis, SypHy ^41^, a recent study showed that glutamate vesicular release on axons starts with myelination onset and occurs at specific heminodal regions coinciding to sites of myelin sheath growth ^25^.

Functional axonal neurotransmission on oligodendroglia would imply the presence of PSD at specific sites along the oligodendroglial cell membrane. PSD are electron dense regions with a high density of neurotransmitter receptors ^31^ that can be visualized at ultrastructural level. We thus searched for the expression of PSD in oligodendroglia, and especially of PSD-95, one of the major canonical scaffolding components of neuronal glutamatergic synapses. OPCs receive glutamatergic-mediated synaptic inputs and express both NMDA and AMPA receptors ^10,11,19,21^ and interestingly, expression of PSD-95 in oligodendroglia was previously reported in OPCs and mOLs in several transcriptomic studies ^32,42–44^. PSD-95 was also detected at the protein level in OPCs ^39^ and along myelin sheath internodes, where it accumulates in paranodal regions ^45^. Herein, we showed that PSD-95 is located in discrete clusters along OPC and mOL membranes. The level and pattern of expression was maintained throughout differentiation, indicating that axon-OL synaptic like interactions are maintained beyond the OPC stage. Strikingly in the zebrafish larva, we found that oligodendroglial PSD-95 clusters are closely apposed to axonal domains, coinciding to sites of glutamatergic synaptic vesicles accumulation. Overall, our data indicate that fundamental structural components of synapses are maintained between axons and mOLs and even increased in number during myelin sheath elongation.

To decipher the role of PSD-95 in oligodendroglia, we generated a cell-specific Gal4 mediated CRISPR-Cas9 knock-out of *dlg4* in the zebrafish and assessed its impact on myelination. As PSD-95 is highly expressed in neurons and other CNS cell types ^32,39,44,45^, CRISPR-Cas9 mediated deletion of *dlg4* allowed us to assess accurately the function of this protein in oligodendroglia, thus avoiding potential confounding effects of this deletion in other cells. We demonstrated that OPC-specific loss-of-function of *dlg4a*, using the *Tg(Olig1:KalTA4)* ^32^ driver line, impacted severely OL maturation, as Olig1-expressing cells exhibited a ramified immature OL morphology in *dlg4* KO with respect to control. As the *Tg(Olig1:KalTA4)* line does not allow the fate mapping of *dlg4* KO cells past the OPC stage, we thus expressed our *dlg4* CRISPR construct in the *Tg(mbp:Gal4)* ^30^ strain to target PSD-95 deletion in mOLs. Importantly, PSD-95 loss-of-function in mOLs leads to shorter myelin internodes, without affecting the number of internodes per cells. Interestingly, our data, in agreement with the recent study of Li and colleagues ^39^, demonstrate that oligodendroglia PSD-95 is required for axon-mOL neurotransmission that regulates oligodendrocyte maturation and myelination in an activity-dependent manner. While Li et al.^39^ generated a double *dlg4a/b* KO, in our present study, we found that *dlg4a* loss-of-function alone in oligodendroglia impedes myelin sheath growth.

PSD-95 clusters and anchors ions channels, glutamatergic receptors and trans-synaptic adhesion proteins at excitatory neuronal synapses ^31^. One could postulate that PSD-95 has the same binding partners in oligodendroglia, as RNA-sequencing data also revealed the expression of PSD-95 binding partners in OPCs and mOLs ^43,44^. PSD-95 is closely opposed to pre-synaptic proteins, such as synapsin and synaptophysin ^39^, and recent *in vivo* studies in zebrafish showed the presence of active axonal vesicular release ^25^ at the elongation sites of myelin internodes. PSD proteins, like gephyrin which is specific of inhibitory synapses, were also observed at calcium activity hotspots in oligodendroglia ^39^, indicating that activity-dependent vesicular release of neurotransmitters increases calcium transients in oligodendroglia ^23^. These calcium transients play a key role in myelin sheath elongation, stability and retraction ^23,24^. Based on these findings, PSD-95 downstream signaling pathways may regulate calcium signaling that mediates the local translation of *mbp* transcripts ^2,36^, and thus timely and spatially restricted myelin sheath growth along electrically active axons. Our data also support the idea that electrical activity could continuously shape myelin microstructure and remodeling, that could have profound impacts on neuronal circuit functions ^5,46^. There has been considerable progress during the last decades in unraveling functional roles of myelin in neuronal circuits that allow complex behaviors ^6–8^ or that even play a role in their dysregulation ^47^. However, the mechanisms by which electrical activity shapes myelination remain so far largely unexplored. Our study will hopefully contribute to better insights into the mechanisms of activity-mediated myelination that has broad implications in neurological diseases.

### Limitations of the study

OLs sense neuronal activity through axon-glial interactions, comprising both functional synapses ^1,2^ and extra-synaptic modes of communications ^48,50^. However, the functional role of axon to oligodendroglial neurotransmission is still poorly understood. Here, we demonstrated that oligodendroglial cells express post-synaptic density proteins and establish synaptic-like microdomains of communications with electrically active axons, which are required for myelin sheath growth. Our study provides deeper insights into mechanisms mediating neuronal activity dependent myelin formation and plasticity. While we provided strong evidence indicating that OLs express PSD-95, the ultrastructure localization of this protein in myelin internodes remain to define accurately. Moreover, PSD-95 binding partners, such neuroligin and AMPA and NMDA receptors were not examined in our study. Further analysis should help to determine whether PSD-95 partners are also necessary for myelin sheath formation and growth. It also remains to elucidate if oligodendroglia PSD-95 function in myelination is required only for myelination of glutamatergic neurons or has a broader role in CNS myelination.

To assess whether PSD-95 microdomains are local hotspots of crosstalk signaling between axons and OLs, electrophysiological recordings of mOLs are required. However, these experiments are technically challenging, due the low capacitance and resistance of myelin sheaths ^52^. These experiments would also require recordings of nanoscale currents in restricted subdomains of mOLs. Recordings of local calcium transients in mOLs, at specific contact sites with axons, could be a valuable proxy to decipher whether PSD-95 microdomains elicit intracellular signals in OLs and how those regulate myelin gene expression.

## Supporting information

Supplemental video S1. Calcium transients in OPCs, related to Figure 1K.

Supplemental video S2. Calcium transients mOL exhibits, related to Figure 1L

Supplemental video S3. Olig1+ cells in control spinal cord at 5 dpf, related to Figure 3D.

Supplemental video S4. Olig1+ cells in dlg4a KO spinal cord at 5 dpf, related to Figure 3D.

## Acknowledgements

We thank Dr Claire Wyart (Paris Brain Institute - ICM Paris) for fruitful comments and feedbacks during this study, and for kindly providing *Tg(s1020t:Gal4)* fish line ^35^. We are grateful to Sophie Nunes, Antoine Arneau, Nolwenn Jezequel (PHENO-Zfish facility, ICM) for fish care, and the ICM imaging facility (ICM Quant) for confocal imaging and electron microscopy. We thank Prof. Koichi Kawakami (NIG, Japan) for sharing the Tol2 system plasmids; Dr. Max Suster for sharing the *Tg(UAS:BoTxBLC-GFP)* transgenic line; Pr. David A. Lyons (University of Edinburgh, UK) for the *Tg(mbp:EGFP)* line and *sox10:GCaMP6* construct; Dr. Bruce Appel (University of Colorado, USA) for *Tg(sox10:mRFP)*, *Tg(Olig2:dsRed)* and *Tg(Olig2:GFP)* lines; Pr. Tim Czopka (University of Edinburgh, UK) for the *Tg(Olig1:KalTA4)* line; Dr. James Jontes (The Ohio State University, USA) for the UAS:synaptophysin-GFP construct; Dr. Martin P.Meyer (King’s College London, UK) for the UAS:PSD-95/GFP construct; Dr. Mariano Soiza-Reilly (Sorbonne University, France) for advices on pre- and post-synaptic antibodies. This study was supported by grants from ANR (RPV14017DDA), the program “Investissements d’Avenir” ANR-10-IAIHU-06 (IHU-A-ICM), ANR-11-INBS-0011 (NeurATRIS) and the French MS Foundation, France Sclérose en Plaques. M.A.M. was supported by a PhD fellowship from Sorbonne university and NeurATRIS, M.G. was funded by postdoctoral fellowships from France Sclérose en Plaques and IBRO.

## Author contributions

B.N.O. supervised the study, designed experiments, and prepared the manuscript. M.A.M. and M.G. designed experiments, generated reporter constructs and lines, performed experiments and prepared the manuscript. M.P. carried out experiments.

## Declaration of Interests

The authors declare no competing interests.

## Supplemental figures and legends

Video S1. Calcium transients in OPCs, related to Figure 1K.

Video S2. Calcium transients mOL exhibits, related to Figure 1L.

Video S3. Olig1+ cells in control spinal cord at 5 dpf, related to Figure 3D.

Video S4. Olig1+ cells in *dlg4a* KO spinal cord at 5 dpf, related to Figure 3D.

## STAR★Methods

### Key resources table

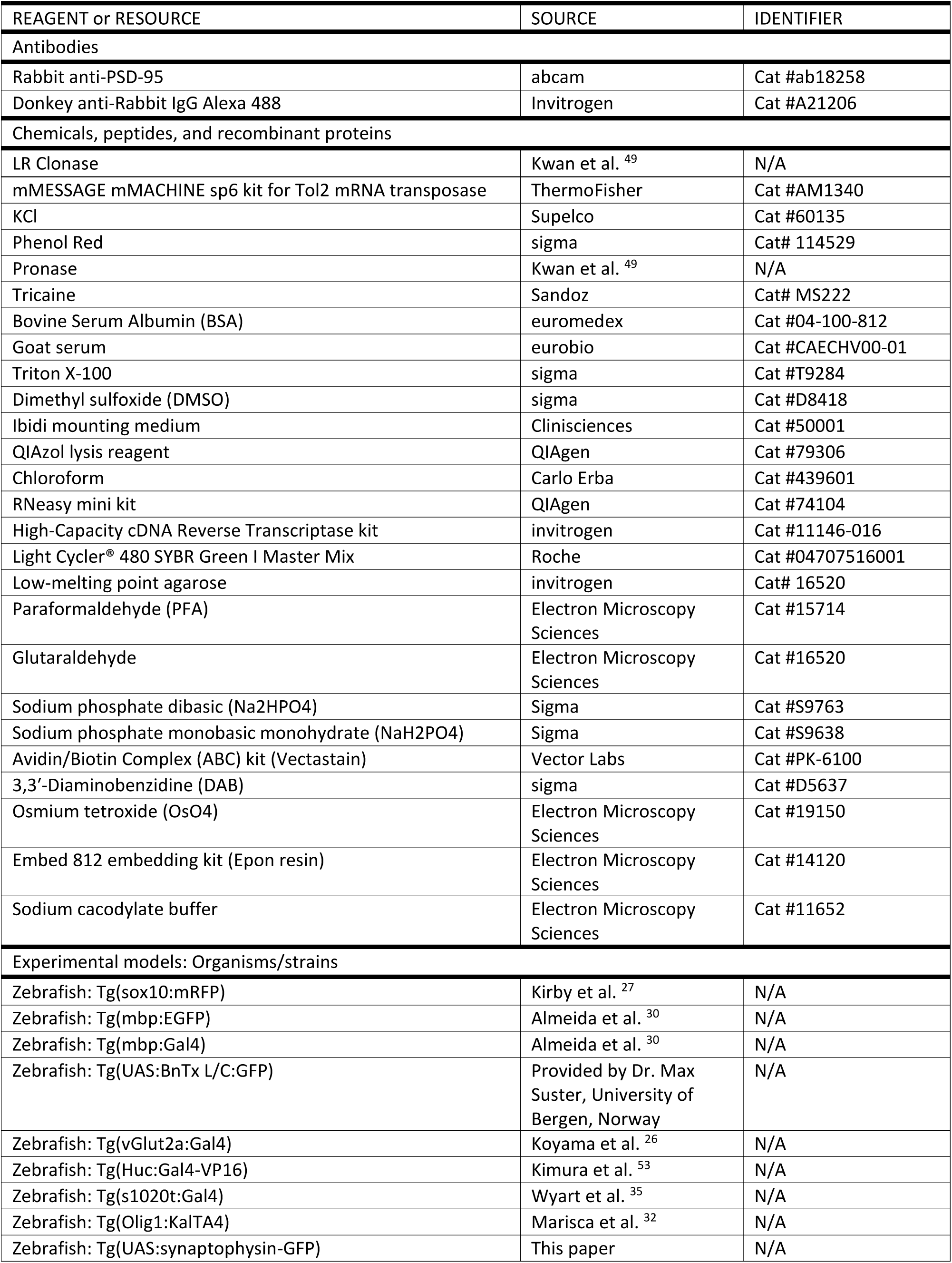
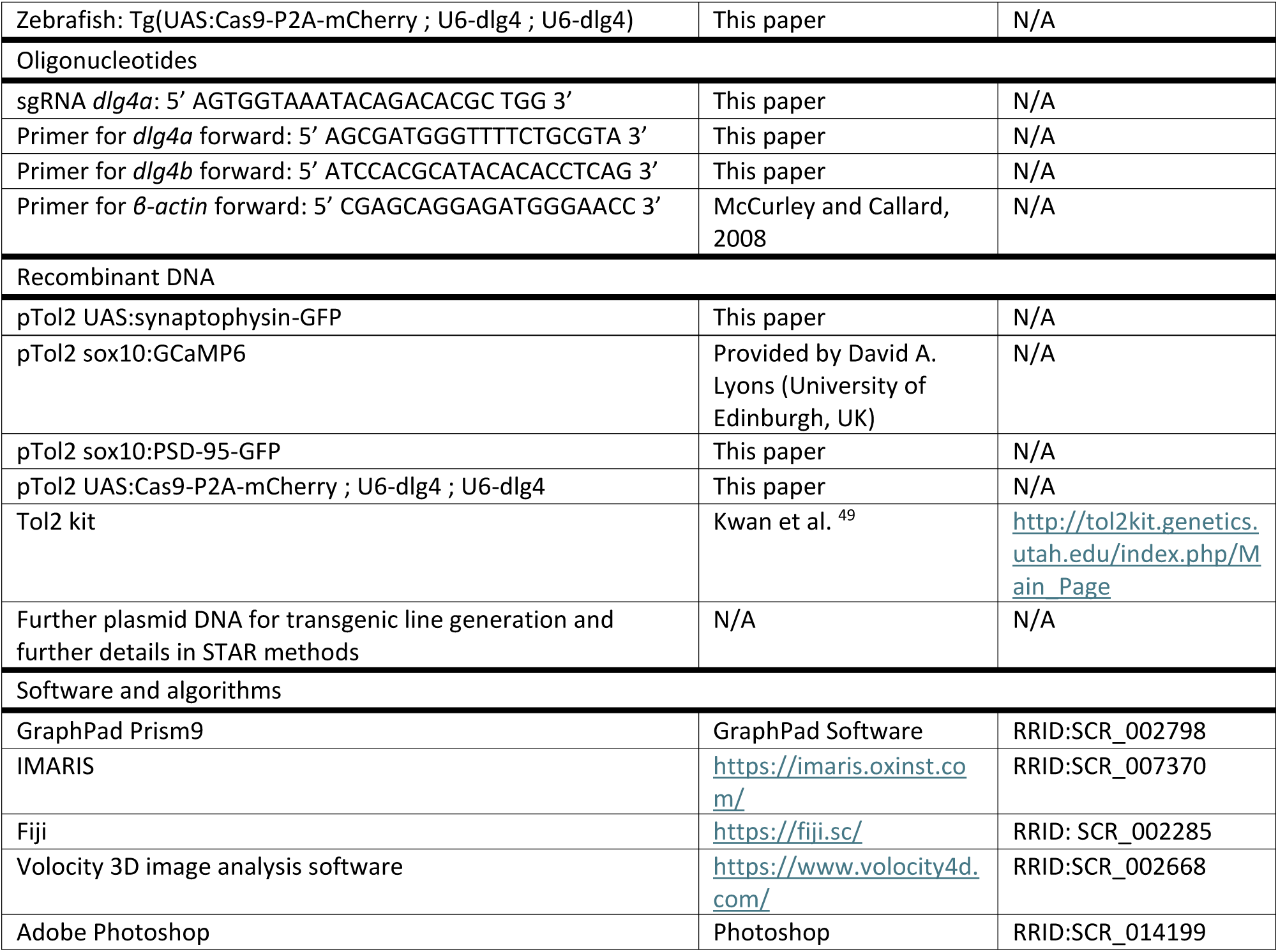

### Resource availability

#### Lead contact

Further information and requests for resources and reagents should be directed to and will be fulfilled by the lead contact, Brahim Nait-Oumesmar.

#### Materials availability

Zebrafish transgenic lines and plasmids generated in this study are available upon request directed to the lead contact, Brahim Nait-Oumesmar.

#### Data and code availability

- Microscopy data reported in this paper will be shared by the lead contact upon request.
- This paper does not report original code.
- Any additional information required for the analysis of data reported in this work paper is available from the lead contact upon request.

### Experimental models and study participant details

#### Zebrafish lines and maintenance

Zebrafish (*Danio rerio*) were maintained on a 14/10h light cycle and water was regulated at 28.5°C, conductivity at 500 μS and pH at 7.4 (Westerfield, 2000). *Tg(sox10:mRFP)*^27^ line were provided by Dr. Bruce Appel, University of Colorado, USA. *Tg(mbp:EGFP)* and *Tg(mbp:Gal4)*^30^ were provided by Pr. David Lyons, Centre for Discovery Brain Sciences University of Edinburgh, UK. *Tg(UAS:BnTx L/C:GFP)* was provided by Dr. Max Suster, University of Bergen, Norway. *Tg(vGlut2a:Gal4)*^26^ was provided by Dr. Joe Fetcho, Cornell University, USA. *Tg(Huc:Gal4-VP16)* ^28^ was obtained from Dr. Shin-ichi Higashijima, National Institute for Physiological Sciences, Japan. *Tg(s1020t:Gal4)* ^35^ was provided by Dr. Claire Wyart, Paris Brain Institute, France. *Tg(Olig1:KalTA4)* ^32^ was provided by Pr. Tim Czopka, Center for Clinical Brain Sciences, University of Edinburgh, UK. *Tg(UAS:synaptophysin-GFP)* and *Tg(UAS:Cas9-P2A-mCherry; U6-dlg4; U6-dlg4)* were generated as described below. Embryos were staged according to Kimmel et al. ^54^. Ages are expressed as day post-fertilization (dpf). Animal and cell numbers are indicated in figure legends. All procedures were approved by the Institutional Ethics Committee of the Institut du Cerveau - ICM, Paris, France.

### Method details

#### Generation of expression plasmid constructs

To generate the *UAS:synaptophysin-GFP* tol2-flanked plasmid, *UAS:synaptophysin-GFP* construct (kindly provided by Dr James D. Jontes; Ohio State University, USA) was subcloned into a Gateway pEntry1A vector and recombined with 5’ (p5E-MCS, Tol2kit, Dr Chien laboratory, University of Utah, USA) and 3’ (p3E-polyA, Tol2kit) entry vectors together with a destination vector containing Tol2 flanking sites (pDestTol2pA2, Dr. Kawakami’s laboratory, Mishima, Japan) using LR clonase (Kwan et al., 2007). To generate the *sox10:PSD-95-GFP* tol2-flanked plasmid, the UAS:PSD-95-GFP construct (Niell et al., 2004; provided by Dr. Martin Meyer, King’s College London, UK) was subcloned into a pEntry1A vector and recombined with the p5E-p7.2sox10:Gal4 (Dr. Thomas Carney, Institute of Molecular and Cell Biology, Singapore) and p3E-polyA vectors, together with the pDestTol2pA2 vector.

To generate *UAS:Cas9-P2A-mCherry; U6-dlg4; U6-dlg4* tol2-flanked cell-specific CRISPR-Cas9 plasmid, we started from the commercially available plasmid *pUAS:Cas9-T2A-GFP*; U6-sgRNA1; U6-sgRNA2 (Cat #74009, Addgene) generated by Dr F. Del Bene ^55^. T2A-GFP was then replaced by P2A-mCherry-CAAX sequence (Cat #74794, Addgene) using AgeI and XbaI restriction enzymes and the Gibson assembly system. CRISPOR.org ^56^ web tool (http://crispor.tefor.net/, developed by tefor infrastructure) was used to identify our *dlg4a* single-guide RNA (sgRNA). As *dlg4a* genes has five variants in zebrafish, we decided to target exon 5, the first common exon of all isoforms, with sgRNA 5’ – AGTGGTAAATACAGACACGC TGG – 3’. The selection of sgRNA was based on the MIT specificity score (>90) ^33^ and Cas9 predicted efficiency (>50) ^34^. We also choose sgRNA with a minimum number of off-targets. BsaI and BsmBI restriction enzymes were used to insert twice our selected *dlg4a* sgRNA in *UAS:Cas9-P2A-mCherry; U6-sgRNA1; U6-sgRNA2* Tol2 expression plasmid.

#### Micro-injections

One-cell stage embryos were collected and injected with 1 nL of the injection solution containing 25 ng/µL of DNA, 50 ng/µL of Tol2 mRNA transposase (mMESSAGE mMACHINE Sp6 kit, AM1340, ThermoFisher), 2.5 M KCl (60135, Supelco) and 0.02% Phenol Red (114529, Sigma) in MilliQ water.

Embryos were maintained at 28.5°C and screened as soon as 2/3 dpf for transgene expression under a Leica fluorescent stereomicroscope. The *UAS:synaptophysin-GFP* positive embryos in a *Tg(HuC:gal4)* background, and the *UAS:Cas9-P2A-mCherry; U6-dlg4; U6-dlg4* positive embryos in *Tg (s1020t:Gal4); Tg(Olig1:KalTA4)* or *Tg(mbp:Gal4)* were raised until adulthood to establish stable lines. F1 progenies were then screened for germline transmission. CRISPR progenies were examined for oligodendroglial cell morphology and myelin sheaths formation and elongation.

#### Mosaic fish generation

Cell transplantation was performed from *sox10:mRFP* to *HuC:gal4;UAS:synaptophysin-GFP* blastulae. Briefly, donor and receptor eggs were harvested separately and dechorionated with pronase 3% in a glass Petri dish. The embryos were transferred to agar molds (PT-1, Adaptive Science Tools) using a glass Pasteur pipette with a fire-polished tip. 30 to 50 cells were taken from the donor embryos and introduced into the receptor embryos using a capillary needle with a 40 µm diameter bevelled tip attached to a manual injector (CellTram Oil, Eppendorf, Germany). Finally, embryos were transferred to glass Petri dishes and maintained at 28.5°C.

#### Immunostaining

Whole larvae were euthanized with 0.2% Tricaine (MS222, Sandoz, Levallois-Perret, France) and then fixed overnight in 4% PFA (PFA, 1514-S, Electron Microscopy Sciences). After several washing steps with 0.1 M PBS, heads and yolks were removed to allow penetration of antibodies. Larve were then incubated overnight in blocking solution (2 mg/mL of BSA, 10% goat serum, 0.5% Triton X-100 (T9284, Sigma), 1% DMSO in 0.1M PBS). Anti-PSD-95 antibody (ab18258, abcam, rabbit IgG, 1,1000 dilution) was incubated 2 days in 2 mg/mL of BSA, 10% NGS, 0.7% Triton X-100, 1% DMSO in 0.1M PBS buffer.

After several washing steps, spinal cords were incubated overnight with rabbit anti-IgG conjugated Alexa 488 (A21206, Invitrogen, 1/1000 dilution) in 1 mg/mL BSA, 1% NGS, 0.5% Triton X-100, 1% DMSO in 0.1M PBS. Spinal cords were mounted with ibidi mounting medium (50001, Clinisciences).

#### Zebrafish dissociation and RT-qPCR

Whole larvae were euthanized with 0.2% Tricaine (MS222, Sandoz, Levallois-Perret, France) and dissociated into QIAzol lysis reagent (79306, QIAgen). Chloroform addition allowed to retrieve RNA that is finally precipitated with isopropanol. RNA extraction was then performed using the RNeasy Mini Kit (74104, QIAgen) according to manufacturer’s instructions. Total RNA harvested was assessed using Nanodrop (Applied Biosystems). cDNA was then synthesized using the High-Capacity cDNA Reverse Transcriptase kit (11146-016, Invitrogen). Finally, qPCR reaction was performed using Light Cycler® 480 SYBR Green I Master Mix (04707516001, Roche) and specific probes for *dlg4a* (5’ AGCGATGGGTTTTCTGCGTA 3’; 5’ GCGAACTCTCCACCAGTAGT 3’) and *dlg4b* (5’ ATCCACGCATACACACCTCAG 3’; 5’ CAACATCTCCGTCCATACCGT 3’). β-actin was used as housekeeping gene. The ΔΔCt method was used to calculate mRNA abundancy and mRNA fold changes.

#### Confocal imaging

Larvae were anesthetized in 0.02% Tricaïne (MS222, Sandoz, Levallois-Perret, France) and mounted with 1.5% low-melting point agarose (16520, invitrogen).

To study pre-synaptic puncta onto oligodendroglial cells and glutamatergic connectivity, spinal cords were imaged using an Olympus FV-1000 confocal microscope with a 40X water immersion objective using 473 and 543nm laser lines. Z-stacks (0.8µm z-step) were taken and whenever applicable, 3D reconstructions were performed using the Volocity software (Perkin Elmer, USA). Pre-synaptic synaptophysin:GFP puncta (larger than 0.35 μm in diameter, Niell et al., 2004) in close apposition with individual sox10:mRFP oligodendroglial cells were visualized and quantified.

To study PSD-95 puncta in oligodendroglia and oligodendroglia-specific *dlg4* loss-of-function, spinal cords were image with a Leica inverted SP8 X white light laser confocal microscope with a 63X oil or a 25X water immersion objective using 488 and 561 nm laser lines. Z-stacks (0.5 µm or 1.02 µm) were taken and Olig1-expressing oligodendroglial cells as well as MBP+ internodes were 3D reconstructed and quantified with IMARIS (Oxford Instruments).

#### Calcium imaging

Three dpf embryos derived from eggs injected with the *sox10:GCaMP6* plasmid (provided by Pr. David Lyons, Centre for Discovery Brain Sciences, University of Edinburgh, UK) were screened for fluorescence. Those with positive cells were selected and cells with oligodendroglial morphology were registered for calcium transients using an Olympus FV-1000 confocal microscope, by z-stacks with a frame interval of 30 seconds. The ΔF/F. of maximal projection images was subsequently analyzed with Image J.

#### Electron microscopy

*Tg(vGlut2a:Gal4;UAS:GFP::sox10:mRFP)* and *Tg(vGlut2a:Gal4;UAS:BnTx/LC:GFP::sox10:mRFP)* 5dpf fish were anesthetized with Tricaine and fixated in 0.1% (16520, Electron Microscopy Sciences) in sodium cacodylate buffer (11652, Electron Microscopy Sciences). After rinsing in 0.1M PB, fish were mounted in agarose and 40μm-longitudinal sections were obtained with a Leica VT1000S Vibratome. Sections were incubated with 2.5% glutaraldehyde (16520, Electron Microscopy Sciences), followed by fixation with 2% osmium tetroxyde (OsO4, 19150, Electron Microscopy Sciences) solution and tissue dehydratation. Next, the tissue was incubated in Epon resine (embed 812 embedding kit, 14120, Electron Microscopy Sciences) at 56°C to allow polymerization. Molds were cut at a UC7 Leica ultramicrotome. Ultrathin sections were observed at the transmission electronic microscope (Hitachi HT7700).

### Quantification and statistical analysis

All graphics and statistical tests were performed using GraphPad Prism9. Shapiro-Wilk normality test was used to assess the Gaussian distribution of the data. Statistical tests were done depending on the type of distribution. Two-tailed unpaired Student’s t test was performed for normally distributed groups, while non-normally distributed groups were compared using the Mann-Whitney test. Statistical tests used are indicated in legends with the number of fish, cells or internodes studied. All data are represented as mean ± standard deviation excepted for Figure 3 where data are represented as violin plots. p-values are indicated on the figure legends, with * p < 0.05; ** p < 0.01, *** p < 0.001; **** p < 0.0001.

## Notes

### Competing Interest Statement

The authors have declared no competing interest.

